# Sub-millisecond optogenetic control of neuronal firing with two-photon holographic photoactivation of Chronos

**DOI:** 10.1101/062182

**Authors:** E. Ronzitti, R. Conti, E. Papagiakoumou, D. Tanese, V. Zampini, E. Chaigneau, A.J. Foust, N. Klapoetke, E.S. Boyden, V. Emiliani

## Abstract

Optogenetic neuronal network manipulation promises to at last unravel a long-standing mystery in neuroscience: how does microcircuit activity causally relate to behavioral and pathological states? The challenge to evoke spikes with high spatial and temporal complexity necessitates further joint development of light-delivery approaches and custom opsins. Two-photon scanning and parallel illumination strategies applied to ChR2- and C1V1-expressing neurons demonstrated reliable, in-depth generation of action potentials both *in-vitro* and *in-vivo*, but thus far lack the temporal precision necessary to induce precisely timed spiking events. Here, we show that efficient current integration enabled by two-photon holographic amplified laser illumination of Chronos, a highly light-sensitive and fast opsin, can evoke spikes with submillisecond precision and repeated firing up to 100 Hz. These results pave the way for optogenetic manipulation with the spatial and temporal sophistication necessary to mimic natural microcircuit activity.

## INTRODUCTION

Optogenetic manipulation of neuronal excitation and inhibition (Boyden et al., 2005; Yizhar et al., 2011a) has the potential to exponentially increase neuroscience’s capacity to elucidate the causal relationships between neural circuits and behavior (O’Connor et al., 2009). Reaching this goal depends on the capability of mimicking physiological neuronal network activity, which ultimately includes optical control of spiking at high firing rates of single selected neurons or specific subsets of neurons (Lin, 2011; Peron and Svoboda, 2011). Progressing in this direction requires combining the engineering of fast and highly light-sensitive opsins with the development of sophisticated light delivery approaches.

Recently engineered channelrhodopsin variants (Berndt et al., 2011; Gunaydin et al., 2010; Klapoetke et al., 2014) exhibit the fast kinetics necessary to drive sustained and precisely timed firing (sub-millisecond jitter; firing rate up to 200 Hz) with visible light, single-photon excitation (1PE). However, scattering and lack of optical sectioning inherent to 1PE pose critical limitations to single-cell photostimulation deep in tissue. Typical 1PE photostimulation schemes use wide-field illumination (through light-emitting diodes or optical fibers implanted in the brain) of large extents, activating every opsin-expressing cell in the illuminated volume (Huber et al., 2008; Tsai et al., 2009). Consequently, the photoactivation specificity relies exclusively on genetic-targeting strategies. This restricts the opsin-expression to specific cell-types (Huang and Zeng, 2013; Luo et al., 2008), but lacks precise optical control of genetically identical cell subpopulations. Spatial selectivity can be achieved by pinpointing specified neurons with light-targeting optical methods (Arrenberg et al., 2010; Dhawale et al., 2010; Papagiakoumou, 2013; Reutsky-Gefen et al., 2013; Szabo et al., 2014; Wyart et al., 2009). However, due to visible light scattering, these approaches have limited penetration depth and poor axial confinement.

Light-delivery approaches based on two-photon excitation (2PE) (Denk et al., 1990) assure the necessary micrometric activation confinement and scattering robustness to reach in depth photostimulation of individual neurons *in vitro* and *in vivo* (Bègue et al., 2013; Packer et al., 2015, 2012; Papagiakoumou et al., 2013, 2010; Prakash et al., 2012; Rickgauer and Tank, 2009; Rickgauer et al., 2014). They can be divided in two main groups: scanning and parallel techniques. Scanning methods quickly move a laser beam across several positions and photostimulate *N* cells with a temporal resolution of *T* = *n* · (*t_d_* + *t_s_*) · *N*, where *n* is the number of positions visited within each cell, and *t_d_* and *t_s_* are the dwell and scanning time, respectively. Temporal current integration with serial scanning can be improved by utilizing opsins such as C1V1 (Yizhar et al., 2011b), featuring slow channel turn-off time, τ_*off*_, and nano-Ampere-currents. This enables action potential (AP) generation at firing frequencies up to 40 Hz (Prakash et al., 2012). Although the overall photostimulation time can be shortened by activating multiple cells through multiple beams scanning (Packer et al., 2012), the combination of sequential scanning and slow opsins limits the achievable temporal precision of photostimulation (photostimulation time for single AP generation *in vitro* and *in vivo*: 5-70 ms; latency=18-58 ms; jitter: 6-20 ms) (Packer et al., 2015, 2012; Prakash et al., 2012).

With “parallel” illumination methods, such as computer generated holography (CGH) or generalized phase contrast (GPC), all selected target regions are excited simultaneously. The temporal resolution for photoactivation is determined primarily by actuator kinetics and is independent of the number of targets, that is *T* = *t_d_*. These configurations (Papagiakoumou et al., 2010) have enabled reliable *in vitro* AP generation in single and multiple cells with millisecond temporal resolution, using ChR2 (Papagiakoumou et al., 2010) and C1V1 (Bègue et al., 2013) at spiking rates up to 30 Hz. Notably, efficient current integration achieved with parallel approaches enables cell’s activation with opsins featuring rapid *τ_off_* (Papagiakoumou et al., 2010). This fact suggests that the combination of ultrafast opsins with 2PE parallel photostimulation can provide an optogenetic toolkit for precise spatial and high temporal optical control of neural firing. Here we demonstrate that such degree of precision can be achieved by using 2PE holographic shaped photostimulation with the opsin Chronos (Klapoetke et al., 2014).

Corroborating what has been demonstrated under single photon illumination, Chronos exhibits unprecedented fast on- and off-kinetics under two-photon activation. This property, combined with the efficient current integration achievable under 2PE parallel illumination with amplified laser pulses, enables reliable AP generation with submillisecond temporal precision and neuronal firing at high frequency under very low power densities. Our results highlight the potential of this approach to mimic a broad range of physiological firing patterns with sub-millisecond temporal precision, key to analyzing how specific patterns of network activity contribute to behavior and pathological states.

## RESULTS

### Photophysical properties of Chronos under Two-Photon Illumination

To determine Chronos kinetic parameters and wavelength sensitivity under 2PE, we used Chinese Hamster Ovary (CHO) cultured cells transfected with a plasmid encoding GFP-tagged Chronos. We used extended 2P holographic patterns (Lutz et al., 2008; Papagiakoumou et al., 2008) shaped over the cell’s surface to photoactivate Chronos-expressing cells and whole-cell patch-clamp recordings to measure the light-induced ionic currents (Figure1a). A high-energy pulse, diode-pumped, ultrafast fiber amplifier laser (pulse frequency 500 kHz, pulse width 250 fs, maximum energy per pulse 20 μJ, *λ* =1030 nm) acted as the photostimulation source, allowing illumination at high peak power.

**Figure 1.**
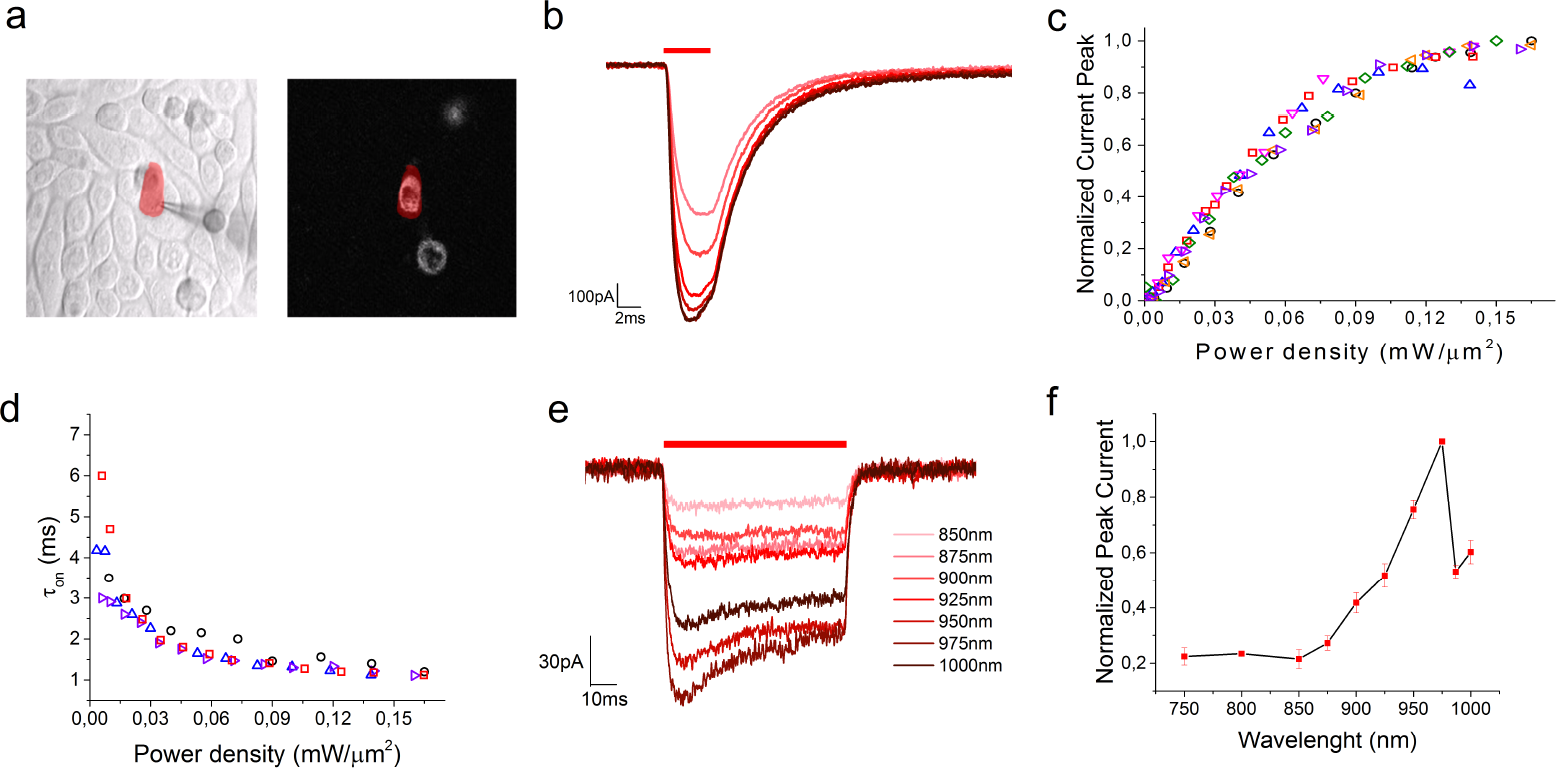
Chronos two-photon kinetics and spectra. (a) IR-transmitted (left) and 2PE fluorescence image (right) of Chronos-expressing CHO cells. A patched cell is photostimulated by a holographic illumination pattern matching its surface (red pattern); (b) Photocurrents generated by irradiating a Chronos-expressing CHO cell with different illumination power densities (P=0.04, 0.06, 0.09, 0.1, 0.12, 0.14 mW/μm^2^; Δt=4 ms; λ=1030nm); (c) Normalized peak photocurrent (n=6 cells) and (d) on-kinetics (measured as time to 90% of the peak) over different irradiation power densities (n=4 cells) (different symbols and colors correspond to different cells); (e) Photocurrents generated by irradiating a Chronos-expressing CHO cell with different illumination wavelengths using equal photon flux; (f) Chronos two-photon action spectra (plotted data are mean ± s.e.m.; n=4 cells).

Initially, we studied the amplitude and kinetics of the photocurrents elicited by photostimulating with light pulses of varying power densities (Figure 1b). Photocurrent’s amplitude increases with the illumination power density up to a saturation regime of the conducting-state (Rickgauer and Tank, 2009) (Figure 1c). As previously reported under 1PE activation (Klapoetke et al., 2014), the temporal dependence of Chronos 2PE-photocurrents reveals unprecedented fast opening and closing channel rates (Figure 1d, Figure Suppl.1). The 2PE turn-on time (*τ_on_*) diminishes by increasing the illumination power (Figure 1d) and reaches a value of 1.2±0.1 ms (n=4 cells) at a power near saturation regime, while *τ_off_* is almost independent on the illumination power and equal to 3.8±0.2 ms (Figure Suppl.1).

We then examined Chronos spectral responses by measuring the amplitude of the photocurrents elicited at various illumination wavelengths (Figure 1e). We used in this case an optical system equipped with a tunable Ti:Sapphire laser as photostimulation source combined with Generalized Phase Contrast (GPC) to generate illumination spots entirely covering the cell (see methods). Photon flux was maintained constant across wavelengths for each cell, ranging between 1.7-2.4×10^26^ photons/m^2^s. Within the range of tunable wavelengths (750-1000 nm), the 2P action spectrum shows an absorption bandwidth between 850 nm and 1000 nm peaking at 975 nm (Figure 1f). A red-shifted shoulder absorption peak is notable at 1000 nm, suggesting that higher absorption rates can be possibly reached at higher wavelengths.

### Two-Photon Chronos spiking

To assess the spiking fidelity of Chronos-expressing neurons under 2-photon illumination in brain slices at various irradiances and frequencies, we injected layer 2/3 of visual cortex of 4 weeks old Swiss mice with adeno-associated virus rAAV8/Synapsin-Chronos-GFP. Slices were prepared 6-8 weeks post-injection. We photoactivated Chronos-expressing neurons by illuminating the soma of the cells with 2PE holographic circular spots (*λ*=1030 nm, fiber amplifier laser system). We recorded cellular activity by whole-cell patch-clamp technique (Figure 2a).

**Figure 2.**
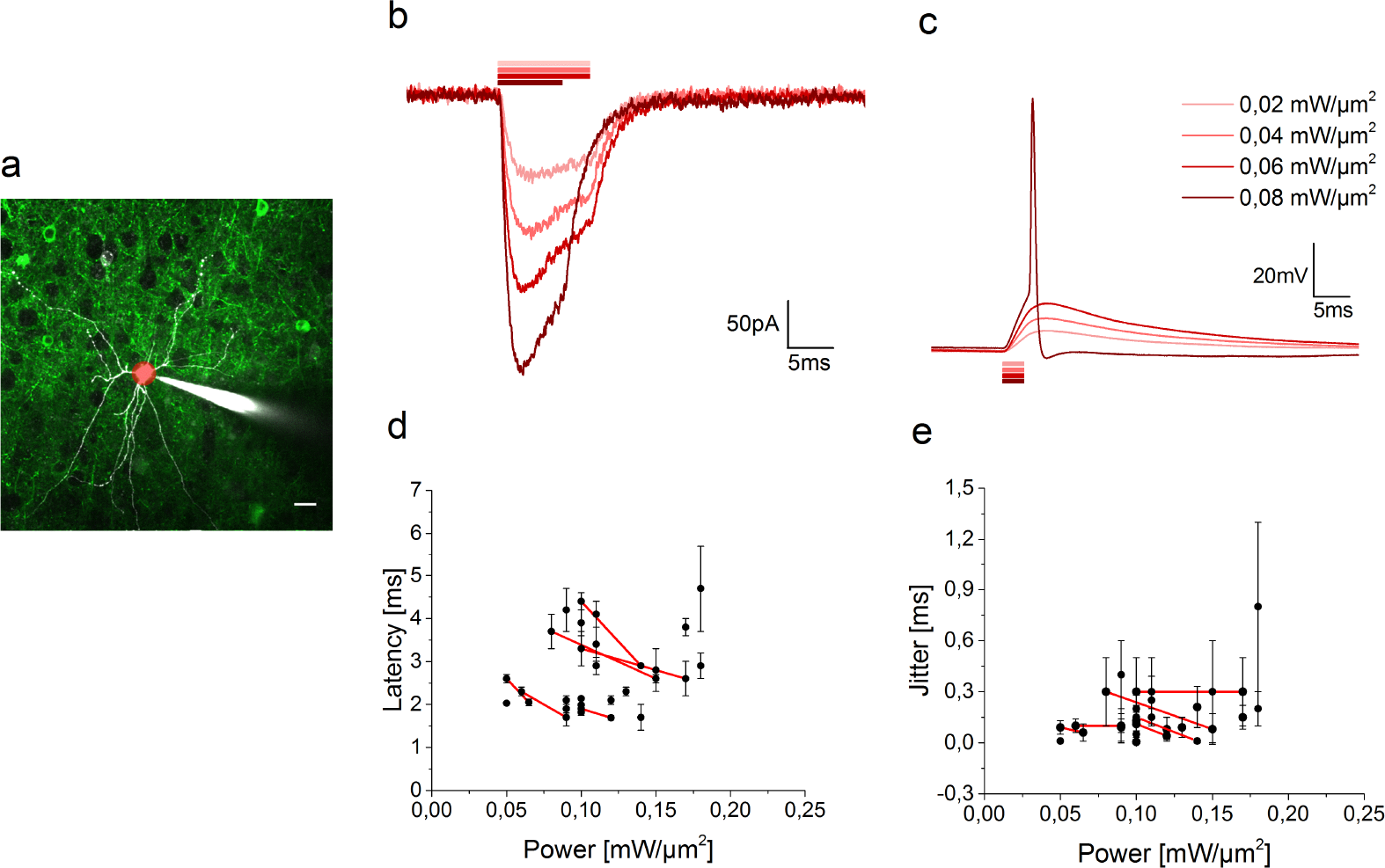
Two-photon activation of Chronos-expressing neurons. (a) Two-photon image of a portion of an acute cortical brain slice of a virus-injected mouse. A patched/Alexa 594-filled (white) interneuron of layer 2/3 expressing Chronos-GFP (green) is photostimulated by a 10-μm holographic spot (red spot). Scale bar 20 μm. (b) Light-evoked photocurrents and (c) membrane potential depolarization over different illumination powers. (d) Latencies (calculated from the start of the light pulse to the spike depolarization threshold) and (e) jitter (calculated as mean deviation of latency) of light-evoked spikes (n=24 cells) by 10-15 μm diameter spots (plotted data are mean ± s.d.). Red lines connect data from same cells.

To demonstrate the achievable temporal precision at high firing rates, we mainly focused the analysis and recordings on fast spiking cells (interneuron); nevertheless, firing properties of slow spiking cells (pyramidal) were also investigated (Table Suppl.1, Figure Suppl.2).

In voltage clamp, photocurrents of hundreds of pico-amperes were evoked upon photostimulation at moderate power density levels (*I*=415±213 pA, *P*=0.08±0.02 mW/μm^2^; mean ± s.d. on *n*=38 interneurons) (Figure 2b, Figure Suppl.3). In current clamp, reliable generation of single AP was obtained using 2-3 ms irradiation durations (Figure 2c) (cells successfully photoeliciting single AP: 74% for *P*<0.12 mW/μm^2^; 84% for *P*<0.3 mW/μm^2^, *n*=38 interneurons). In a few cases of highly expressing cells (peak photocurrent in voltage-clamp *I*=817±315 pA, *P*=0.08±0.02 mW/μm^2^, n=5 cells) we could shorten the stimulation pulse and still obtain reliable AP generation with 1 ms (n=2 cells) and 0.7 ms (n=3 cells) pulse duration (*P*=0.13±0.05 mW/μm^2^).

Action potentials occurred with short latencies (2.83±0.97 ms, *P*=0.11±0.03 mW/μm^2^; mean ± s.d. on *n* =24 cells, Figure 2d) and sub-millisecond temporal precision (jitter 0.19±0.16 ms, *P*=0.11±0.03 mW/μm^2^; mean ± s.d. on *n*=24 cells, Figure 2e). Increasing the illumination power enabled reducing spike latency and jitter on a cell (see line-connected cells in Figure 2d-e).

The short spike latency, the short pulse duration necessary to trigger an action potential, and the fast turn-off channel kinetics suggest that high frequency spiking can be driven in Chronos-expressing neurons under 2PE holographic-illumination. We thus photostimulated with trains of light pulses (2-3 ms pulse width) at different frequencies neurons that successfully elicited a single AP. We found that 2PE light stimulation could drive cells to spike up to a frequency of 100 Hz (Figure 3a). Reliable 2PE driven spiking (with no failures) was achieved up to a stimulation frequency of 50 Hz, while spike failures might be encountered at 100 Hz stimulation frequency after 3-4 pulses, depending on the Chronos expression level and the cell type (Figure 3b). It is worth mentioning that, contrary to what obtained in slower opsins (Boyden et al., 2005; Gunaydin et al., 2010), we observed small plateau potentials, even at higher frequency, indicating fast repolarization between illumination pulses (Fenno et al., 2011). Furthermore, no asynchronous spikes were evoked and a modest increase of spike latency and jitter throughout the succession of photostimulations was recorded (Figure 3c and Figure 3d). Both observations confirm the high spike temporal precision of Chronos when driven by 2PE patterned light during sustained stimulation of short light pulses. Importantly, thanks to the short pulse duration and relatively small amount of light necessary to trigger an AP, prolonged series of photostimulations did not induce physiologically relevant cellular perturbation. In a series of experiments (n=5 cells) we tested cell stability during several minutes of successive series of ten light-pulses delivered at 10 Hz every 6 s. The AP amplitude, input resistance and membrane potential, did not change appreciably during 40 consecutive train stimulations (average AP amplitude: 99±4 mV; AP amplitude variation: 0.8±0.4 mV; input resistance: 143±17 MΩ; input resistance variation 0±6 % membrane potential variation: 0.9±0.8 mV) (Figure Suppl.4a). Spike latencies showed modest augmentation during successive trains of photostimulation, suggesting that the proportion of inactivated channels gradually incremented during long-term, sustained photostimulation of few minutes (Figure Suppl.4b).

**Figure 3.**
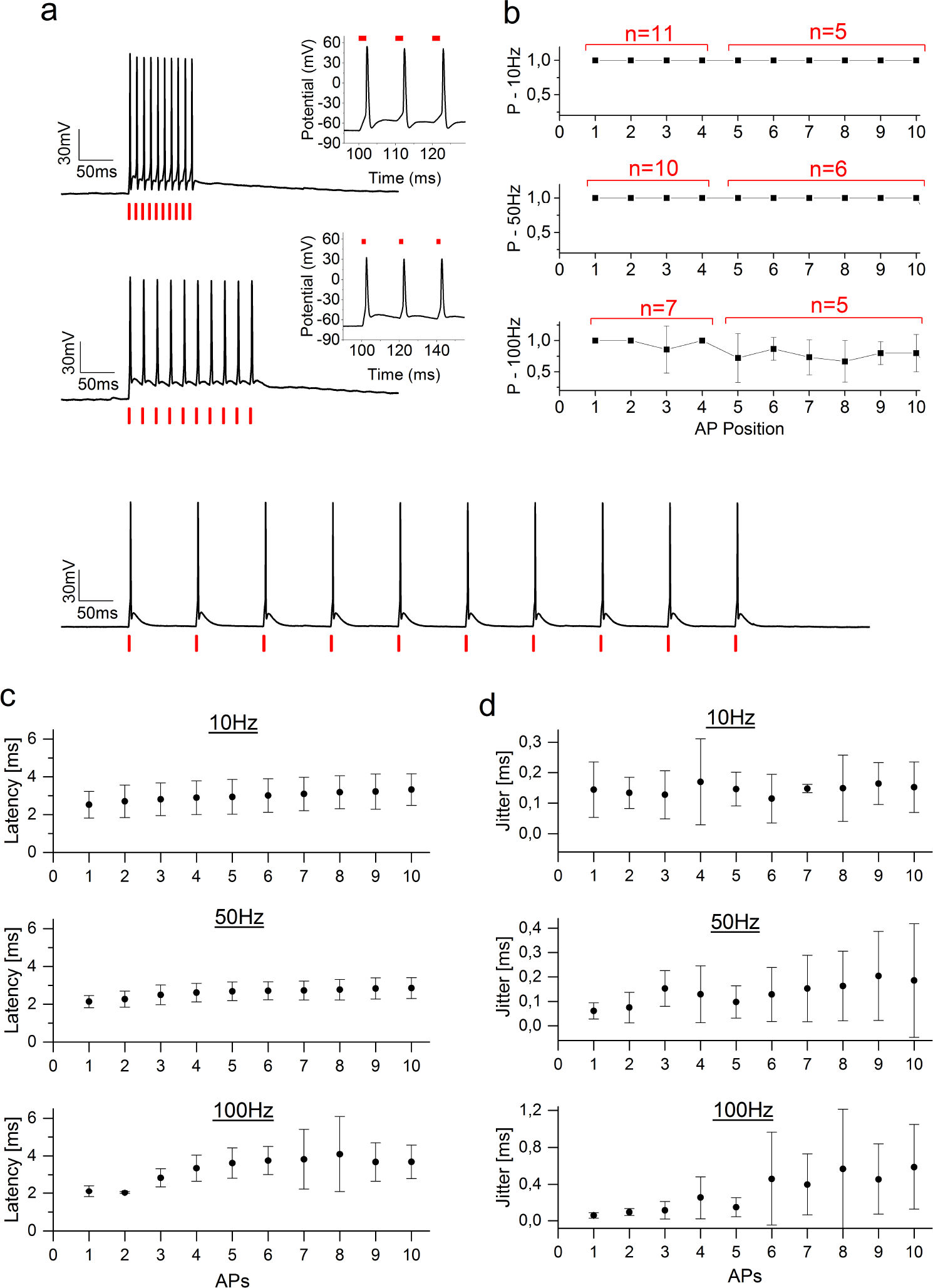
Driven spiking fidelity over different frequencies. (a) Sample trains of 10 photostimulation pulses of 2-ms duration each, over different frequencies (10 Hz, P=0.09 mW/μm^2^; 50 Hz, P=0.09 mW/μm^2^; 100 Hz, P=0.12 mW/μm^2^). (b) Spike probability within the ten pulses (cells have been photostimulated either with four or ten light-pulse trains; in red it is indicated the number of cells where the i-th spike has been tested). (c) Latency and (d) jitter of each light-evoked spike of the train. Illumination power range 0.05-0.17 mW/μm^2^, 2-3 ms illumination pulse duration, 10-15 μm diameter spot, λ=1030 nm. Plotted data are mean ±s.d.

The short off-kinetics of Chronos will probably make it difficult to achieve efficient current integration with serial laser scanning; however, this limitation could turn into an advantage for applications combining optical photostimulation with Ca^2+^ imaging, where the main challenge is to avoid spurious photostimulation during imaging (Emiliani et al., 2015). Indeed, a typical experimental design for 2P Ca^2+^ imaging, uses fast scanning of a diffraction-limited spot. In the case of Chronos, this should generate only weak photocurrents due to the fast channel closing rate, despite the strong spectral overlap of Chronos and calcium indicators of the GCaMP family (Chen et al., 2013), that are commonly used. To test this hypothesis, we performed whole-cell patch-clamp recordings of membrane potential variations in Chronos-expressing neurons during 2PE raster scanning of a 920-nm femtosecond beam typically used for imaging genetically encoded calcium indicators (GCaMPs). In particular, we tested cell depolarization while performing 2P scanning imaging using two FOVs (300 μm and 150 μm) at different frame rates obtained by changing scanning speed and inter-line distance (n=6 cells). While scanning a small FOV (150 μm, P=14 mW) could induce in few cases high cell depolarization (2 out of 6 cells, Figure Suppl.5), increasing FOV to 300μm reduced cell depolarization far below the spiking threshold, independent on the scanning parameters. Using 2PE imaging with resonant scanners or Red Calcium indicators should enable enable reducing depolarization even further

## DISCUSSION

We have demonstrated that the high light sensitivity and fast on-and off-kinetics of Chronos, combined with the efficient current integration of parallel holographic illumination, enable reliable supra-threshold depolarization in neurons of acute cortical slices, with millisecond latencies and submillisecond jitters, under 2-photon excitation.

Millisecond temporal control of neuronal firing opens the door to spatially and temporally mimicking neural activity patterns, critical to studying how spike temporal and rate codes convey information in the brain (Baranauskas, 2015; Izhikevich et al., 2003; Lisman, 1997), refine reverse-engineering neural circuits programs (O’Connor et al., 2009), as well as closed-loop and activity-guided perturbation of neural dynamics and behavior (Grosenick et al., 2015). Precise spike timing and high frequency neural events have been linked to many relevant phenomena in neural plasticity, behavior, and pathology, including setting the direction of synaptic change in spike-timing dependent plasticity (Dan and Poo, 2004), mediating neural coding in systems such as the auditory system (Köppl, 1997), and contributing to functions altered in diseases such as Parkinson’s disease (Foffani et al., 2003). Some cell types such as parvalbumin-positive interneurons fire at high frequencies, and the loss of such neurons may be a contributor to psychiatric illness (Behrens et al., 2007). Fast patterns of neural spiking may also govern whether spikes propagate throughout dendrites, and thus affect localized cellular processes (Buzsáki et al., 1996).

Efficient current integration achievable under 2P parallel photostimulation can also enable millisecond spike generation and sub-millisecond temporal jitter using opsins with very high light activation efficiency but an order of magnitude slower kinetics. However, in this case, the slow off kinetics cause a slow repolarization between pulses with consequent extra spikes and pronounced prolonged plateau depolarization, thus significantly hampering the fidelity of high-frequency light-driven spiking (Chaigneau et al., in preparation). Conversely, here we demonstrated that the fast kinetics of Chronos enabled sub-millisecond timed control of light-driven neural firing up to 100 Hz, with no extra spike and small plateau potential. Both latency and jitter values were impressively well maintained during high frequency (up to 100 Hz) trains.

We expect that performance similar to the one achieved with Chronos and 2PE holography can be reached by alternative parallel patterning approaches as GPC (Papagiakoumou et al., 2010) or low-NA Gaussian beams (Rickgauer et al., 2014) and/or other fast opsins, as ChETA variants (Gunaydin et al., 2010), although in this last case the inferior light sensitivity (Lin, 2011) will probably require using higher excitation power levels.

Using a fiber-amplifier laser source in our case, enabled higher 2PE efficiency compared to typical mode-locked Ti:Sapphire laser oscillators due to lower pulse repetition rates of the femtosecond beam (two-photon excited signal, 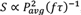, with *f* repetition rate, *τ* pulse width and *P_avg_* average beam power (Xu and Webb, 1996; Zipfel et al., 2003)). This enabled efficient photo-current and spike generation with low light powers (< 0.15 mW/μm^2^; 12-27 mW/cell) and short light pulses (<3 ms) allowing repetitive cell stimulation without affecting cells’ physiological parameters. These findings, together with the fact that amplified lasers can deliver high average power beams (≈10 W at laser output), indicate that laser power is not the limiting factor for the maximum achievable number of targets using CGH. That will be more likely limited by other factors, such as sample heating and deterioration of the photostimulation spatial resolution. Indeed, for multiple-cell stimulation, the photostimulation of cellular processes crossing the illumination volume will ultimately limit the maximum number of targets that can be stimulated by maintaining cellular resolution.

Finally, we have shown that thanks to the fast kinetics of Chronos, imaging under conditions similar to what is typically adopted for GCaMP 2PE imaging (large filed of view, fast and low-resolution scanning) ensure modest membrane potential depolarization. This will enable combining Chronos photostimulation with functional imaging using genetic Ca^2+^ indicators, despite the close spectral overlap between Chronos and GCaMP. Future developments of Chronos viral vector using a red fluorescent reporter will enable to directly prove this assumption. Further reduction of membrane depolarizations during functional imaging could be achieved by minimizing the imaging beam dwell time on the cell using fast raster resonant scanning configurations or smart targeted-path scanning schemes (Grewe et al., 2011; Lillis et al., 2008) to arbitrarily define line-scan trajectories over preselected cells able to minimize the imaging beam dwell time on the cells.

## ACKNOWLEDGEMENTS

We thank Marta Gajowa and Fabio Simony-Conesa for performing part of mice viral injections; Cecile Jouffret for help in cell culture maintenance; Oscar Hernandez for contributing simulation tools. This work received financial support from the Agence Nationale de la Recherche (grants ANR-12-BSV5-0011-01, ANR-12-BSV5-0011, ANR-14-CE13-0016), the National Institutes of Health (NIH 1-U01-NS090501-01), the Medical Research foundation FRM (DVS20131228920), the FRC and the Rotary Club through the program « Espoir en Tête 2012 », and a National Science Foundation International Postdoctoral Research Fellowship (A.J.F.).

## MATERIALS AND METHODS

### Imaging and Photostimulation Setup

The optical system was built around a commercial upright microscope (SliceScope, Scientifica) and combined a multi-light path imaging architecture with a holographic-based photoactivation apparatus.

The imaging system was composed of three different imaging pathways: a two-photon (2PE) raster scanning, a single-photon (1PE) wide-field epi-fluorescence and an infrared (IR) off-center oblique illumination imaging (Figure Suppl.6).

2PE imaging was provided by a mode-locked Ti-Sapphire source (Coherent Chameleon Vision II, pulse width 140 fs, tuning range 680-1080 nm). The femtosecond pulsed beam was raster scanned on the sample via a pair of XY galvanometric mirrors (3 mm aperture, 6215H series, Cambridge Technology) imaged at the back aperture of the microscope objective (40X W APO NIR, Nikon) through an afocal telescope (scan lens: *f*=100 mm, Thorlabs #AC508-300-B; tube lens: *f*=300 mm, Thorlabs #AC508-100-B). Galvanometric mirrors were driven by two servo drivers (MicroMax series 671, Cambridge Technology) controlled by a Digital/Analog converter board (PCI-6110, National Instrument). Emitted fluorescence was collected by a fiber-coupled detection scheme(Ducros et al., 2011) including a large diameter collector lens (*f*=75 mm, Thorlabs #LB1309-A) and a 5-mm-diameter liquid light guide (LLG, Series 300, Lumatec customized with a *f*=14.5 mm doublet lens glued at the fiber entrance by Till Photonics and an anti-reflective coating provided at the fiber exit). The fiber exit was imaged onto two photomultiplier tubes GaAsP (H10770-40 SEL, Hamamatsu H10770-40 SEL) by a set of three matching aspheres lenses (*f*=23.5 mm, Melles Griot #LAG-32.5-23.5-C). Following the fiber exit, fluorescence light was filtered by two infrared-light blocking filters (FF01-750sp, Semrock), split into two channels by a dichroic mirror (FF555-Di03, Semrock) and detected through two emission filters (FF01-510/84 and FF02-617/73, Semrock). 2PE imaging laser power was tuned by combining an electrically-controlled liquid crystal variable phase retarder (LRC-200-IR1, Meadowlark Optics) and a polarizer cube (BB-050-IR1, Meadowlark Optics).

1PE imaging was provided by a LED source (M470L2, Thorlabs) filtered by a bandwidth excitation filter (FF01-452/45, Semrock) and coupled to a diffuser (DG10-1500, Thorlabs) and an achromatic lens (*f*=30 mm, #LA1805 Thorlabs) to provide widefield illumination on the sample. 1PE excited fluorescence was collected through a tube lens (f=200 mm), separated from 1PE excitation light using a dichroic mirror (FF510-Di02, Semrock) and detected by a CCD camera (0rca-05G, Hamamatsu) after passing through a visible bandwidth filter (FF01-609/181, Semrock). Fluorescence induced either by 2PE raster scanning or 1PE widefield illumination was collected by PMTs or CCD respectively, having in the detection pathway a switchable dichroic mirror (FF705-Di01, 70x50mm, Semrock) downstream and a dichroic mirror (ZT670rdc-xxrxt, Chroma) upstream of the fluorescence detection pathway.

The collection of the fluorescence induced by 2PE raster scanning or 1PE widefield illumination through either on PMTs or CCD respectively, is ensured by the presence in the fluorescence detection pathway of a downstream movable dichroic mirror (FF705-Di01, 70x50mm custom size, Semrock) and an upstream dichroic mirror (ZT670rdc-xxrxt, Chroma).

IR oblique illumination imaging was provided by an IR-LED source (M780L2, Thorlabs) installed at the rear port of the microscope coupled with a condenser focusing the light on the sample. IR light transmitted through the sample was collected with an IR CCD (IR-1000, DAGE-MIT).

2PE photoactivation was performed by generating arbitrary light shapes selectively matching specific opsin-tagged patterns in the sample via a spatially-controlled phase modulation of the illumination beam wavefront. In particular, a femtosecond pulsed beam delivered by a diode pumped, fiber amplifier system (Satsuma HP, Amplitude Systemes; pulse width 250 fs, tunable repetition rate 500-2000 kHz, gated from single shot up to 2000 kHz with an external modulator, maximum pulse energy 20 μJ maximum average power 10 W, wavelength *λ*=1030 nm) operated at 500 kHz, was widened through an expanding telescope and reflected on the sensitive area of a reconfigurable liquid crystal on silicon spatial light modulator (LCOS-SLM, X10468-07, Hamamatsu Photonics). The reflected beam was projected at the back focal plane of the objective with an afocal telescope (*f*=300 mm, Thorlabs #AC508-300-B and f=500 mm Thorlabs #AC508-500-B). The SLM was controlled by a custom-designed software(Lutz et al., 2008) based on a Gerchberg and Saxton iterative algorithm(Gerchberg and Saxton, 1972), which converts an arbitrary intensity pattern on the sample plane to a specific phase profile to be addressed at the SLM plane. The effect of the zero-order of diffraction in the sample was suppressed by introducing a cylindrical lens (Hernandez et al., 2014).

2PE imaging scan and 2P photoactivation beams were combined through a large dichroic mirror (T970dcspxr, 50x70 mm, Chroma).

Intensity patterns matching the cellular surface were used for photoactivation of CHO cells, while holographich circular spots of 10-20 μm were used to photoactivate neurons. Power densities in the text are given at the output of the objective and calculated referring to the excitation spot’s surface.

### Cell line culture and electrophysiology

Hamster Chinese Ovary’s cells were cultured in an incubator at 32 °C and 5% CO_2_ in a D-MEM/F12 GlutaMAX medium (life technologies) with the addition of 1mM glutamine, 1% streptomycin and 10% fetal bovine serum. Cells were plated on Thermanox plastic coverslips (Thermo Fisher Scientific, NY, USA) 24 hours prior to transfection. Cells were transfected with a home-made plasmid (FCK-Gene90-GFP provided by Ed Boyden’s laboratory) containing the Chronos opsin bound to GFP. The DNA was transfected using the ExGen 500 (Euromedex,France) transfection reagent and cells were recorded 24-48 hours after transfection. Cultured cells were transferred for recording in a chamber mounted on the headstage of the microscope.Recordings were performed at room temperature. The extracellular medium during electrical recording was of the following composition (in mM): 140 NaCl, 5 KCl, 2 CaCl_2_, 1 MgCl_2_, 20 Hepes, 25 Glucose. PH adjusted to 7.5 by NaOH. Electrophysiological recordings were performed through a Multiclamp 700B Amplifier (Molecular Devices, USA) in the whole-cell voltage clamp recording configuration. Patch pipettes, pulled from borosilicate glass capillaries, had a resistance in the bath that ranged from 4 to 6 MΩs. Intracellular solution was composed of (in mM): 140 KCl, 2 MgCl_2_, 2 Mg ATP, 0.4 Na GTP, 10 Hepes, 20 BAPTA. PH adjusted to 7.3; osmolarity 330. Cells were maintained at −40 mV throughout recordings. The holding current ranged from 0.8 to 80 pA.

### Stereotaxic injections of viral vector

All experimental procedures were approved by the Paris Descartes Ethics Committee for Animal Research (registered number CEEA34.EV.118.12) and followed European Union and Institutional guidelines for laboratory animals care and use (Council directive 86/609 EEC). Stereotaxic injections of the viral vector rAAV8/Synapsin-ChR90-GFP (University of North Carolina vector core) were performed in 4 weeks old swiss mice (Janvier lab). Mice were anesthetized with ketamine (80mg/Kg-xylazine (5mg/Kg) solution and a small craniotomy (0.7 mm) was made on the skull overlying V1 cortex. Injection of 1-1.5 μl solution containing the viral vector was made with a cannula at about 80-100 nl/min at 200-250 μm below the dural surface. The craniotomy and the skull were sutured and the mouse recovered from anesthesia. The combination of our injection protocol and the viral vector used, resulted in a higher percentage of opsin-expressing interneurons with respect to pyramidal cells. Moreover in the latter the level of expression was relatively low as can be seen from the data in Supplementary Table1. In pyramidal cells we in fact had to double the illuminated surface (surface diameter going from 10-15um for interneurons to 15-20um for pyramidal cells) to obtain similar average photocurrents.

### Acute slice preparation and electrophysiology

Acute parasaggital slices of the visual cortex were made from adult mice 6-8 weeks after viral injection. Animals were decapitated after being anesthetized with isofluorane. The brain was quickly removed and put in an ice-cold sucrose cutting solution (in mM: 85 NaCl, 65 sucrose, 2.5 KCl, 0.5 CaCl_2_, 4 MgCl_2_, 25 glucose, pH 7.4 saturated with 95% O_2_ and 5% CO_2_).

The brain was dissected and 300 μm-thick slices were then cut on a vibratome VT1200S (Leica Biosystems, Germany). Slices were maintained at 32 °C for 30 min in standard ACSF, (sACSF composition in mM: 125 NaCl, 2.5 KCl, 26 NaHCO_3_, 1.25 NaH_2_PO_4_, 1 MgCl_2_, 1.5 CaCl_2_, 25 glucose, 0.5 ascorbic acid) saturated with 95% O_2_ and 5% CO_2_ and then transferred at room temperature in the same solution until use. During the experiments, the slices were kept in a chamber and continuously perfused with fresh sACSF saturated with 95% O_2_ and 5% CO_2_. We chose to patch only cells deeper than 30 μm from the slice surface, as more superficial cells are usually more depolarized and risk to have an important proportion of their processes damaged by the slicing procedure. The pipette solutions contained (in mM): 130 K-gluconate, 7 KCl, 4 MgATP, 0.3 GTP, 10 phosphocreatine-Na, 10 HEPES (pH 7.2 with KOH). Borosilicate glass pipette resistance in the bath: 8 - 12 MΩs. Series resistance ranged between 20 and 40 MΩs (avg: 30±7 MΩs) and were compensated for by 60 to 70%. For interneuron’s recordings, input resistance ranged between 105 and 520 MΩs (avg: 220±100 MΩs), membrane capacitance ranged between 18 and 79 pF (avg: 37±15 pF) and membrane resting potential ranged between −54 and −76mV (avg: −65±5 mV). For pyramidal cell’s recordings input resistance ranged between 172 and 770 MΩs (avg: 400±200 MΩs), membrane capacitance ranged between 42 and 158 pF (avg: 90±40 pF) and membrane resting potential ranged between −70 and −82mV (avg: −76±5 mV).

Neuronal type was established based on cell morphology under IR oblique illumination and their firing properties upon suprathreshold current injections of increasing amplitudes up to 350 pA in the current-clamp recording configuration. Cells were visually inspected prior to patch with a 40X water immersion objective. The characteristic AP firing was the main parameter used during recordings to distinguish the type of cells. The firing of pyramidal cells shows a classic spike frequency adaptation and the maximal frequency of firing is close to 20 Hz; the second AP in a series is wider than the first, and the development of an adaptive hump as the cell is further depolarized can be often observed. Most interneurons were fast spiking, which show almost no adaptation in their firing, their AP characterized by a big component of the fast after-hyperpolarization and with frequencies well beyond 20 Hz(Cauli et al., 2000). A few cases (n=3) were regular spiking non pyramidal or stuttering interneurons.

### Photostimulation procedure

Two different approaches were used to search for cells expressing the Chronos opsin. In most of the experiments expression was sparse and it was then relatively simple to detect single transfected cells through wide-field fluorescence illumination. In cases of higher expression, in which the background fluorescence was too high, we used the 2PE imaging. In regions of really high levels of expression we instead adopted the option of blindly patch cells, and their level of expression was verified by electrical recordings of the currents evoked by photostimulation. Recorded cells were monitored through IR oblique light illumination during the patching phase.

#### Photostimulation of CHO cells

2PE photocurrents were evoked while maintaining the cell in voltage-clamp configuration at −40 mV. For the characterization of the current response as a function of the light power density, a holographic shape completely covering the cell surface was used. In this way, all channels fell within the illuminated area and we could avoid problems of spurious recruitment of channels outside of the holographic pattern as the power increases.

The experiments to characterize the spectral response of Chronos were performed on a different setup, equipped with a tunable Ti:Sapphire laser and with a GPC holographic illumination delivering large diameter spots that covered the whole cell.

#### Photostimulation of neuronal cells

Recorded cells were photostimulated with 2PE holographic spots of 10 to 20 μm diameter, centered on the soma of the cell. The duration and intensity of the stimulus varied depending on the experimental procedure and are indicated in the results. The value of the power density used during the stimulation was calculated as the ratio of the laser power measured after the objective and the surface of the designed spot.

### Data acquisition

For the experiments on spectral characterization, electrical signals were recorded through a Multiclamp 700B patch-clamp amplifier and digitized using a Digidata 1322A interface and pClamp software (Axon Instruments). All other recordings were performed using a Multiclamp 700B patch-clamp amplifier and digitized using a National Instrument data acquisition board (NI USB-6259). NeuroMatic software environment in conjunction with IgorPro software (Wavemetrics) were used for data acquisition and analysis. Results are quoted as mean ± s.d., unless otherwise specified in the text. Voltage-and Current-clamp recordings were filtered at 10 kHz and sampled at 50 kHz.

